# Introducing *circStudio*, a Python package for preprocessing, analyzing and modeling actigraphy data

**DOI:** 10.64898/2026.03.30.711342

**Authors:** Daniel Marques, Carina Fernandes, Cátia Reis, Nuno L. Barbosa-Morais

## Abstract

Actigraphy is a non-invasive and cost-effective method for monitoring behavioral rhythms under real-world conditions by collecting time-resolved measurements of locomotor activity, light exposure, and temperature. Although several open-source packages support specific aspects of actigraphy analysis, aspects such as preprocessing, metric calculation, and mathematical modeling are often distributed across separate software packages, limiting interoperability and increasing programming overhead. Here we introduce *circStudio*, a Python package that unifies actigraphy data processing and mathematical modeling of circadian rhythms within a single framework. Built from the *pyActigraphy* codebase and integrating circadian models from the Arcascope *circadian* package, circStudio provides flexible preprocessing tools, support for multiple actigraphy file formats through adaptor classes, standalone functions for computing commonly used actigraphy metrics, and implementations of several mathematical models of circadian rhythms. The package enables users to move efficiently from raw wearable data to physiologically interpretable circadian outputs. Ultimately, circStudio aims to facilitate reproducible workflows and to provide a flexible foundation for research applications across circadian biology, sleep science, and digital health.

## Introduction

Actigraphy is a non-invasive method that uses wearable sensors to record continuous time series of locomotor activity, light exposure and peripheral temperature, enabling the quantification and inference of sleep-wake patterns under ambulatory conditions (Fekedulegn et al., 2020). In contrast with polysomnography (PSG), the gold standard for assessing sleep but a method that is expensive and often disruptive to normal sleep routines, actigraphy is well suited for large-scale data collection owing to its non-invasive nature and relatively low cost (Fekedulegn et al., 2020). Actigraphy time series can be used to derive commonly reported actigraphy variables or features (Carpenter et al., 2025). These features include interdaily stability (IS), which measures the strength and consistency of the 24-hour rest-activity pattern across days in the actigraphy recording; the sleep regularity index (SRI), which quantifies the similarity of sleep-wake patterns between consecutive days; and the relative amplitude (RA), which contrasts the difference between the 10 most active (or illuminated) hours and the least active (or illuminated) hours of the day (Carpenter et al., 2025).

Actigraphy time series can also be used to model the response of the circadian system to a given pattern of light exposure through mathematical models of circadian rhythms (Hannay & Moreno, 2020). One of the most widely used formalisms was originally developed by Kronauer based on a Van der Pol (VDP) oscillator and later refined by Forger and Jewett (Forger et al., 1999; Jewett et al., 1999). More recently, Hannay proposed a formalism aimed at improving the physiological interpretability of model state equations (Hannay et al., 2019). In particular, the Hannay framework describes how the dynamics of rhythms at the level of individual cells can give rise to collective rhythm amplitude in the suprachiasmatic nucleus (SCN) of the hypothalamus (Hannay et al., 2019). Other models extend the VDP formalism to incorporate non-photic input (St. Hilaire et al., 2007), while others simulate the daily trajectory of melatonin across different physiological compartments, such as the pineal gland, plasma and exogeneous melatonin (Breslow et al., 2013).

Over the years, several actimetry sensor brands have been developed, each producing data in distinct proprietary file formats and typically accompanied by dedicated, closed-source analysis software (Hammad et al., 2021). Although these tools facilitate basic processing of sensor data, they often restrict the range of analytical approaches available to researchers and lead to actigraphy data underutilization (Fekedulegn et al., 2020). Moreover, reliance on device-specific software and closed-source processing pipelines makes it difficult to generalize results across studies (Hammad et al., 2021). To address these limitations, several programming packages have been developed. For instance, in the R programming language, a number of packages have been dedicated to actigraphy data analysis, such as *LightLogR* (Zauner et al., 2025) for the analysis of personal light exposure.

Critically, most analytical functionality for actigraphy data has not been integrated into a single, easy-to-use package that supports preprocessing, analysis, and modeling within a unified framework. In Python, one of the most notable open-source toolboxes developed for actigraphy data analysis is *pyActigraphy* (Hammad et al., 2021), which also has a dedicated module for light exposure analysis called *pyLight* (Hammad et al., 2024). Another Python package, *circadian*, enables the calculation of such models (Tavella et al., 2023), but it is not optimized to handle noisy light schedules and often fails to compute model state variables under more realistic light exposure patterns. Moreover, moving between *pyActigraphy* and circadian typically requires additional programming effort. To bridge these tools and provide an integrated workflow for actigraphy analysis and circadian modeling, we introduce circStudio, a Python framework that combines actigraphy data processing with mathematical modeling of circadian rhythms.

### *circStudio* design and architecture

The *circStudio* package was developed on top of *pyActigraphy*, although it introduces several architectural differences. *circStudio* is organized into two main modules: `io`, which handles file input/output operations and preprocessing, and `analysis`, which provides functionality for data analysis. In contrast, *pyActigraphy* distributes preprocessing (e.g., `io`, `log`, `mask`) and analysis (e.g., `metrics`, `sleep`, `light`, `viz`) functionality across multiple modules. A key motivation for reorganizing the codebase was to reduce code duplication and foster community contributions, while providing greater flexibility for users. One example of this flexibility is the segregation of data analysis functions from the Raw class, which is responsible for reading and preprocessing actigraphy files. This architecture enables users to implement custom preprocessing routines and use *circStudio* exclusively to compute actigraphy-derived metrics and/or simulate mathematical models.

A typical workflow in *circStudio* involves preprocessing, feature extraction, mathematical modeling, and visualization of actigraphy data. Raw recordings are first imported and preprocessed to handle device-specific formats using adaptor classes, masking, and data cleaning. The processed time series can then be used to extract commonly reported actigraphy-derived features, including activity, light exposure, and sleep-related variables. The processed data may also serve as inputs to mathematical models of circadian rhythms that simulate the response of the circadian system to light exposure. Finally, *circStudio* provides functionality to visualize the resulting time series, model outputs, and several actigraphy-derived metrics.

### Installation

The latest version of *circStudio* can be retrieved directly from PyPI using:

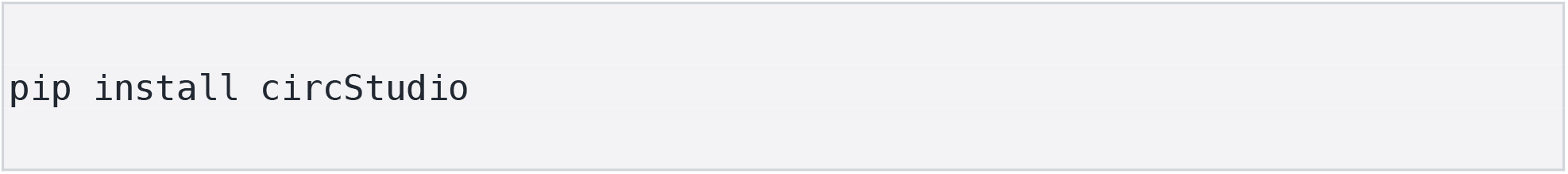

The installation can be verified by running:

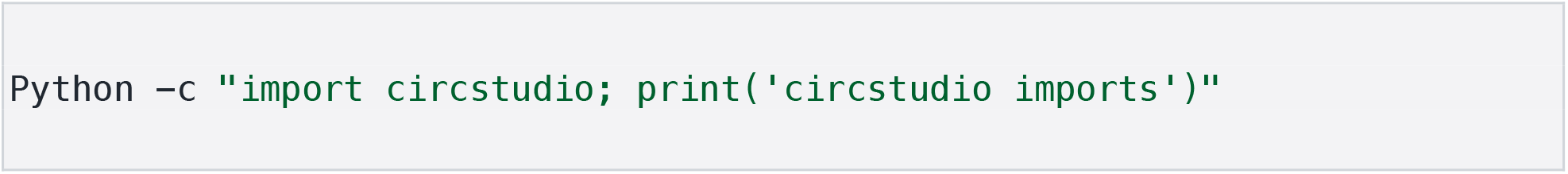

### Example workflows

A typical use case begins by importing the package and loading an actigraphy file, followed by signal preprocessing, metric computation, and optionally simulating circadian system dynamics using a light intensity time series as input to mathematical models of circadian rhythms.

### Loading actigraphy data with the *Raw* class

The first step in any analysis is to import *circStudio*:

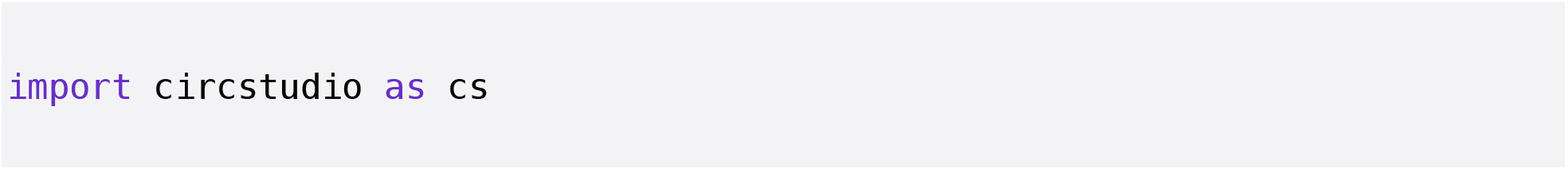

The preferred method for loading actigraphy data in *circStudio* is to create a new instance of the `Raw` class. This requires the actigraphy data to be preformatted as a table containing activity and light series. Unlike *pyActigraphy, circStudio* decouples the metric computation from the `Raw` class, which is used exclusively for signal preprocessing. This design allows users to define custom preprocessing pipelines and to apply circStudio metrics to other types of time series, such as peripheral body temperature.

To create a `Raw` instance, users must specify the dataframe (`df`), the activity (`activity`) and light (`light`) time series (set `light` to `None` if no light time series is available), as well as start time (`start_time`), total duration (`period`), and sampling frequency (`frequency`):

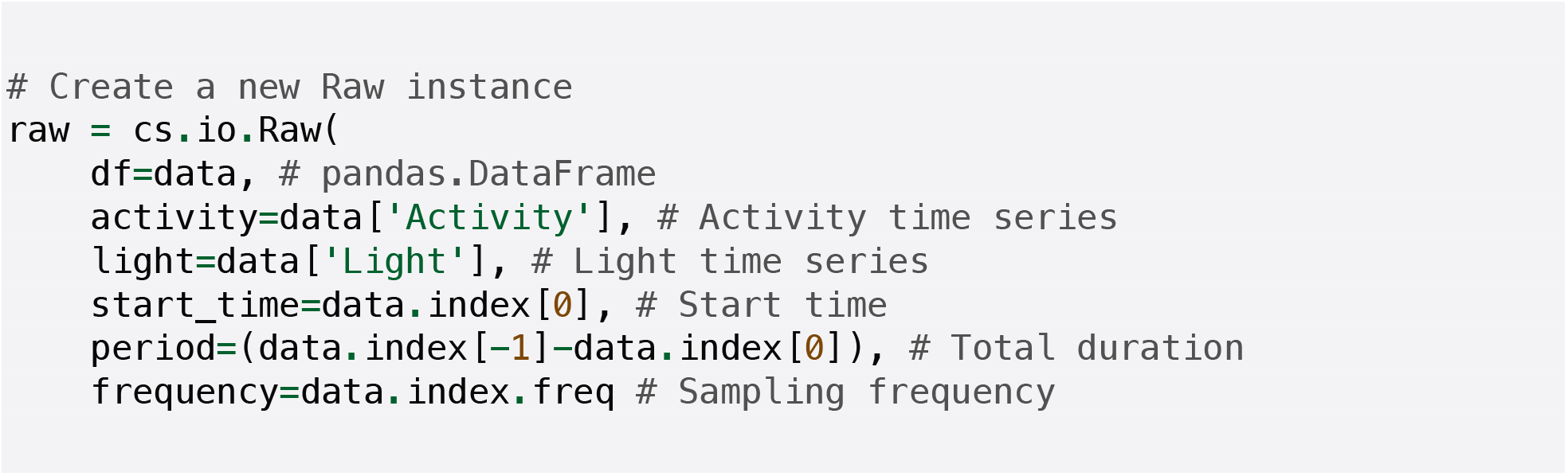

### Reading actigraphy files using adaptor subclasses

Although the `Raw` class is intentionally format-agnostic and flexible, *circStudio* provides several adaptor subclasses for commonly used actigraphy formats. These adaptor classes, migrated from *pyActigraphy*, enable the conversion of supported file types into `Raw` instances with minimal effort. In the following example, a `.txt` file generated by an ActTrust (Condor Instruments) device, included as a sample file, is loaded. To access the sample files distributed with *circStudio*, a file path is first constructed:

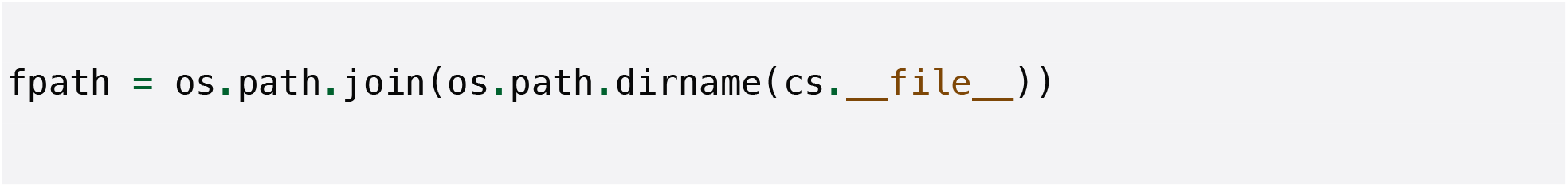

A `Raw` instance can then be created using the helper function `read_atr`:

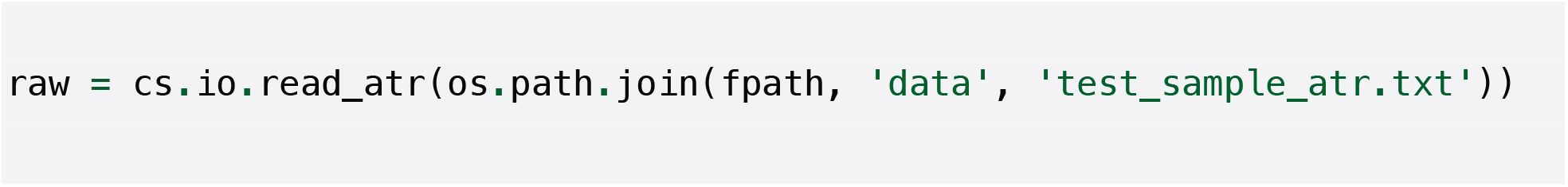

Some ActTrust files contain an extra line above the header, (e.g., *#ActLogModel=2*.*0*.*0*) that should be ignored during import. This can be handled using the `skip_rows` argument:

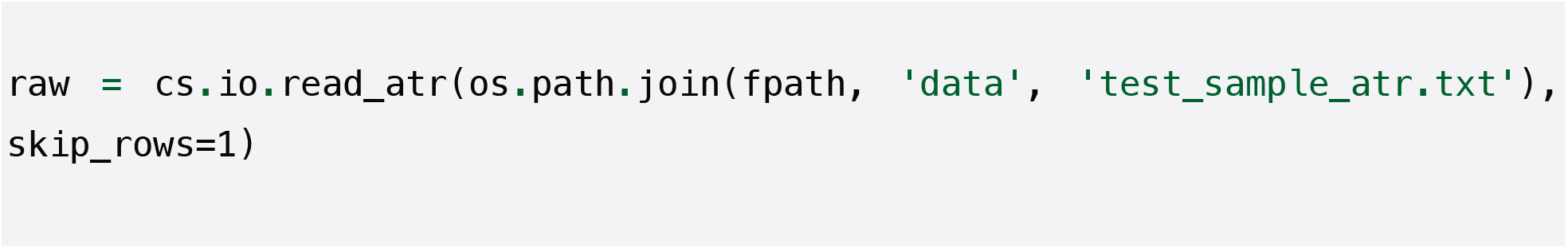

Once an original actigraphy file has been converted into a Raw object, the underlying time series can be accessed directly. For example, the locomotor activity signal is stored as a `pandas.Series` and can be retrieved via `raw.activity`, while the light intensity time series can be accessed via `raw.light`. In both cases, the `plot()`method provides a convenient way to quickly visualize activity and light recordings. For instance, the following command plots the activity time series:

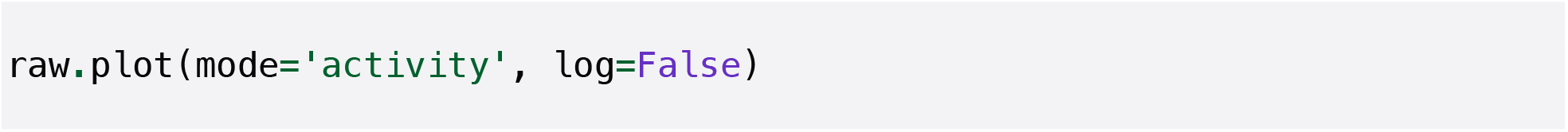

**Figure 1.**
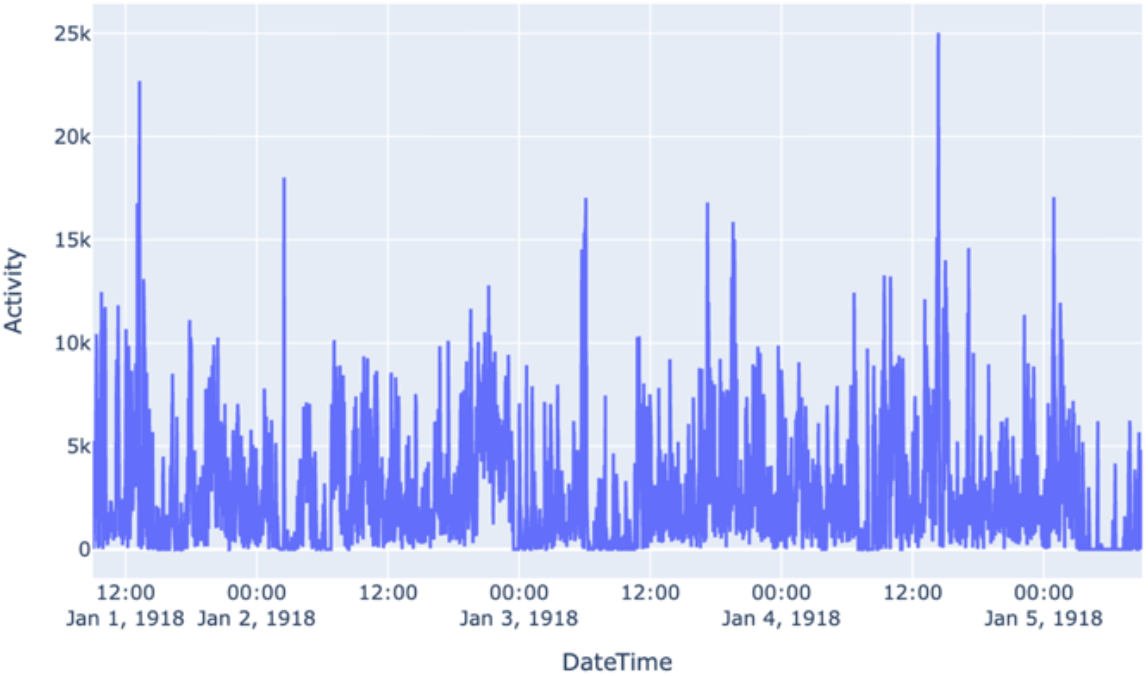
Locomotor activity time series. Calling the *raw*.*plot(mode=‘activity’)* method generates an interactive visualization of the activity signal. To get a similar visualization for the light intensity time series, use *raw*.*plot(mode=‘light’)*.

### Preprocessing actigraphy data for subsequent analyzes

#### Discarding invalid sequences at the start and end of the recording

To remove potentially invalid segments at the beginning and end of a recording, *circStudio* follows an approach similar to that implemented in *pyActigraphy*. Briefly, users can exclude initial portions of the recording by specifying `start_time` and remove terminal portions by defining the duration to retain after `start_time` using the `period` argument:

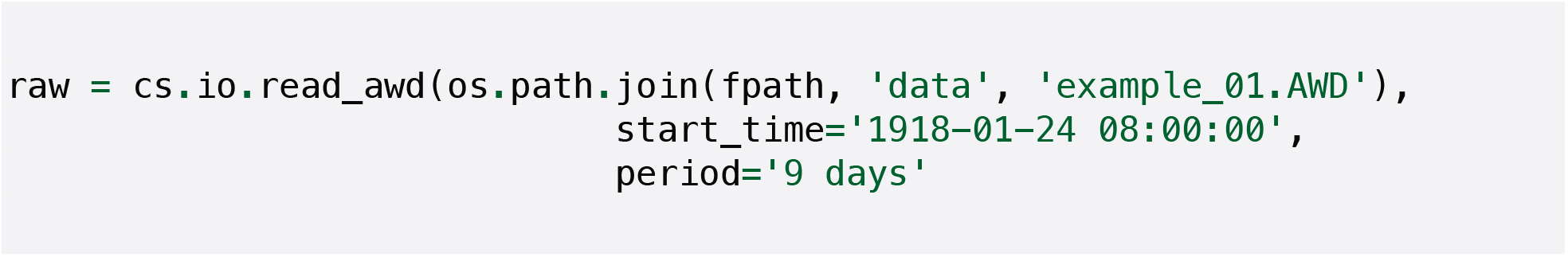

#### Discarding invalid sequences during the recording

*circStudio* allows users to mask spurious periods of inactivity within a recording by marking them as `NaN`. Some downstream analyses, however, require continuous time series, such as fitting a Cosinor model or simulating the response of the circadian system to a given light time series. To address this, *circStudio* provides an imputation procedure that fills missing values using the mean value from the corresponding times across other days in the recording. All masking operations, including data imputation, are handled via the `apply_filters()`method, allowing multiple filters to be applied with a single command.

A mask can be created for inactivity periods lasting at least 120 min:

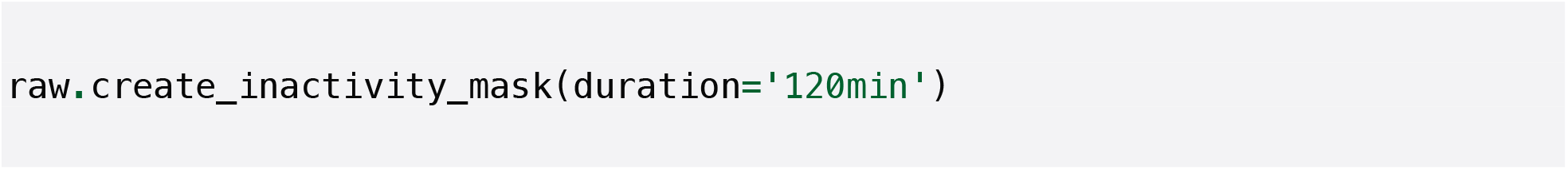

Custom masks can also be added from a CSV file containing *Mask, Start_time, Stop_time*, and an optional label for each inactivity segment to be masked:

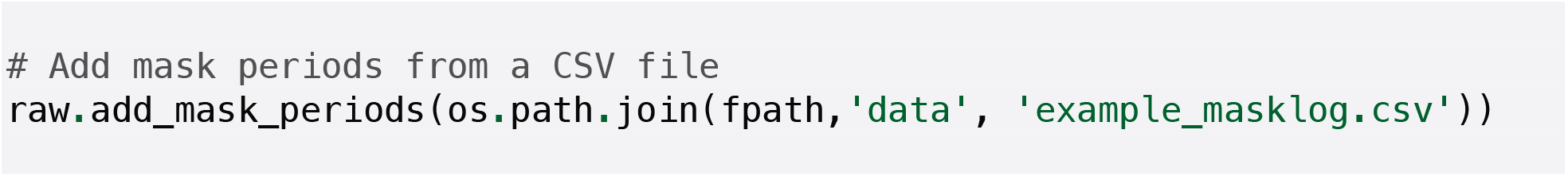

The mask can then be applied and the resulting missing values imputed:

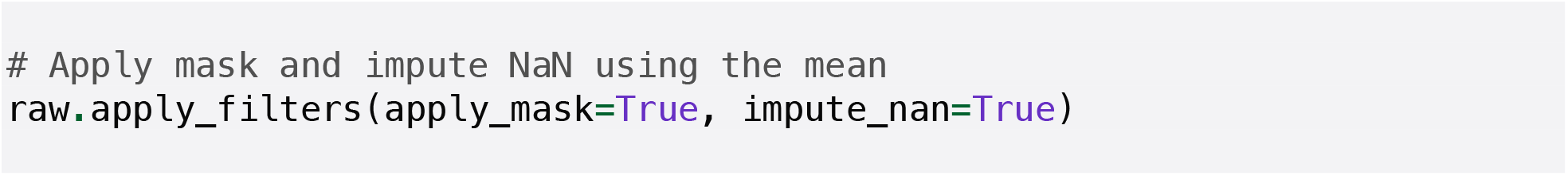

Additionally, `apply_filters()` also enables the signal to be resampled and binarized. For instance, to binarize:

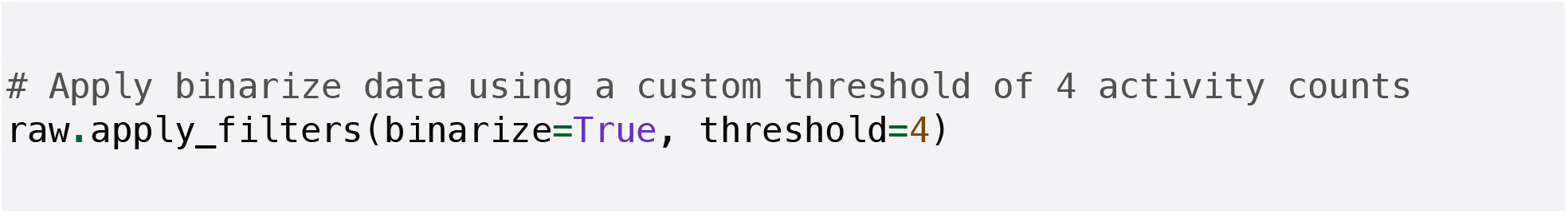

To remove all filters and restore the original signal, the `reset_filters()`method can be used:

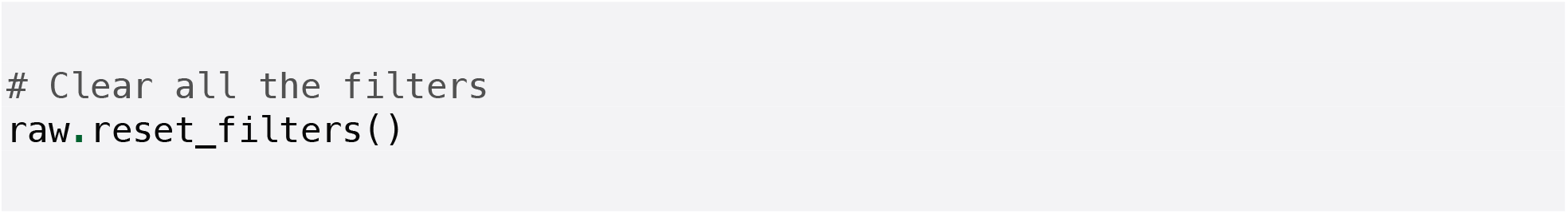

For some applications, such as using mathematical models of circadian rhythms, smoothing may be useful to reduce the effect of extreme fluctuations in the input signal. Thus, *circStudio* provides tools to smooth the activity and light series:

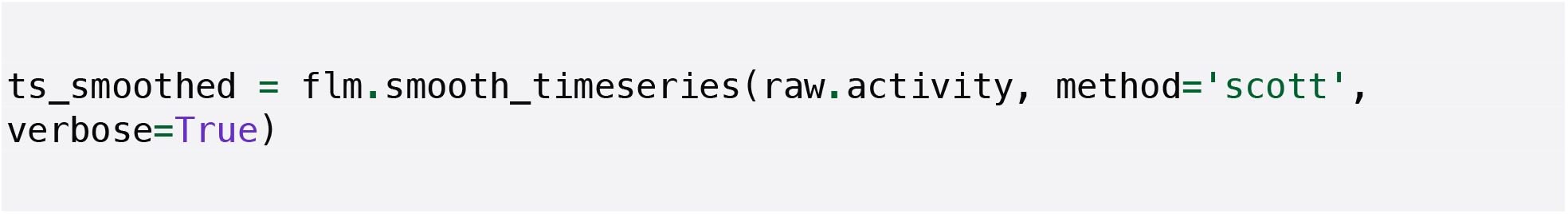

### Computing actigraphy-derived variables

After completing preprocessing, a broad range of actigraphy variables can be calculated. Examples include total average daily activity (ADAT), interdaily stability (IS), and the average activity during the 10 most active hours of the day (M10). In contrast to *pyActigraphy*, metrics are implemented as standalone functions requiring a data argument, which should be a *pandas*.*Series* index by date and time. Of note, the time series used for these calculations should be preprocessed beforehand, for example using `apply_filters()`.

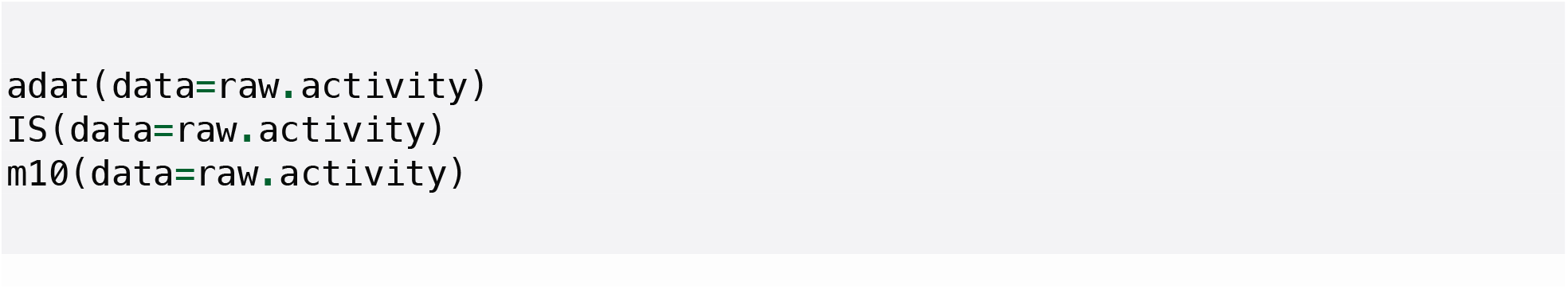

### Simulating mathematical models of circadian rhythms

*circStudio* implements a suite of mathematical models of circadian rhythms developed to predict human circadian physiology: `Forger` (Forger et al., 1999), `Jewett` (Jewett et al., 1999), `HannaySP` and `HannayTP` (Hannay et al., 2019), `Hilaire07` (St. Hilaire et al., 2007), `Breslow13` (Breslow et al., 2013), and `Skeldon23` (Skeldon et al., 2023). The model equations and parameters were retrieved from the circadian package developed by the Arcascope team (Tavella et al., 2023). The general workflow for these models parallels that used for metric computation: a light intensity time series is first imported and preprocessed, after which a model instance is created. For example, a new instance of the Forger model can be initialized using the following syntax:

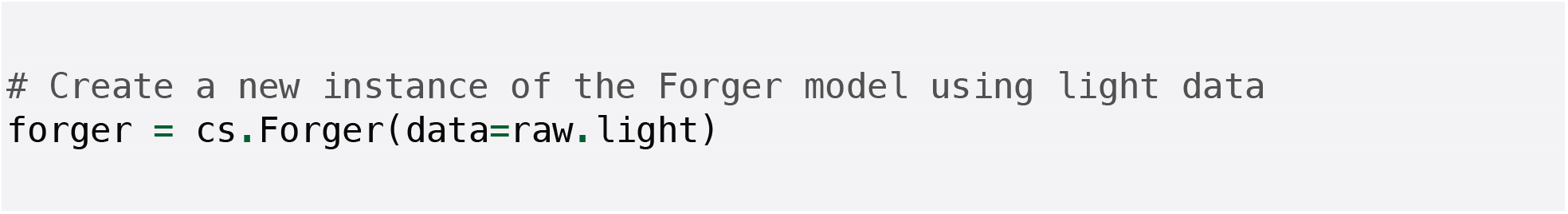

With the exception of `Hilaire07` and `Skeldon23`, model initialization syntax is consistent across implementations:

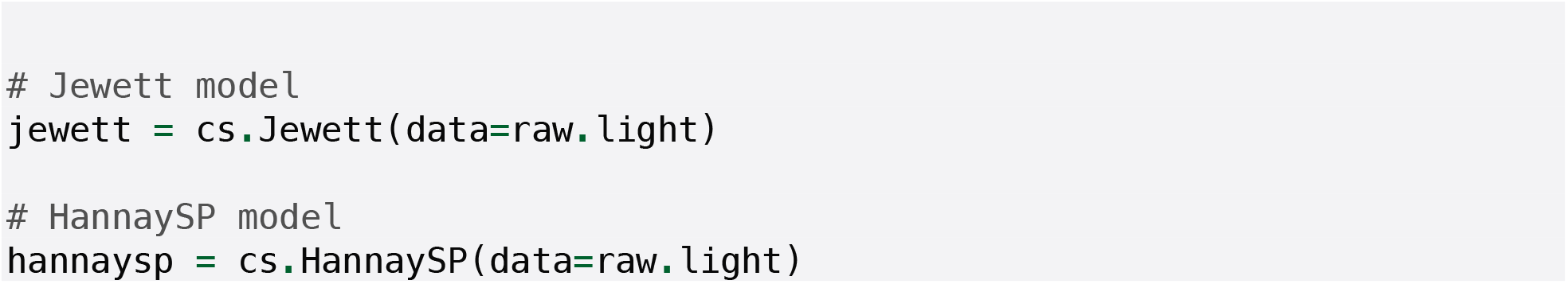

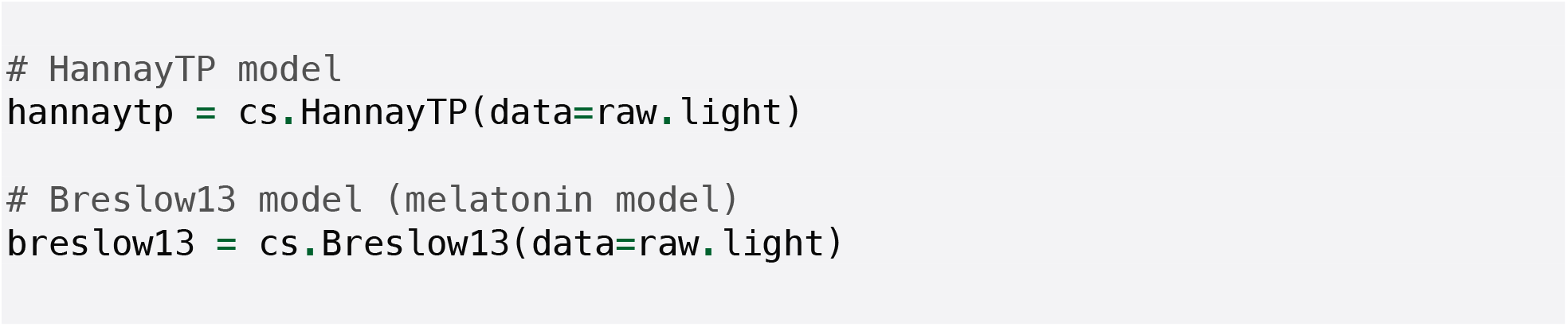

The `Hilaire07` model requires a light time series and a sleep schedule as inputs. By default, the sleep schedule is inferred using the Roenneberg algorithm, although a user-defined schedule may also be provided via the `sleep_algo` argument. Any externally supplied sleep schedule must share the same datetime index as `raw.light`. This argument can also be used to select alternative sleep algorithms, such as Crespo:

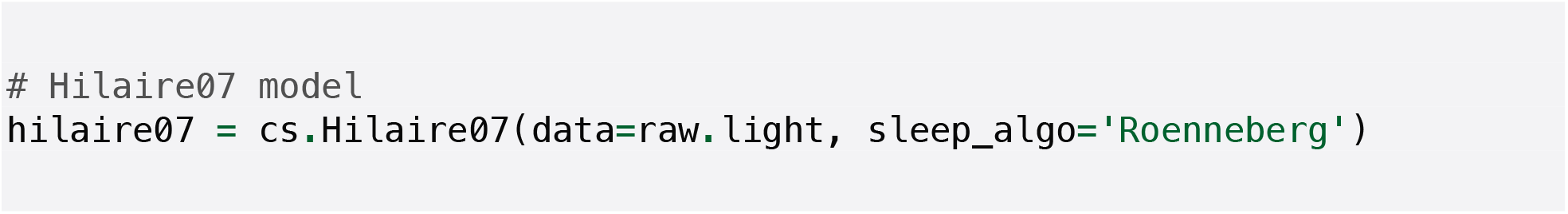

Before extracting phase markers, it is often useful to estimate new initial conditions from the observed light time series. Because these models are formulated as systems of differential equations, their numerical solution depends on the initial state. circStudio provides default initial conditions to allow immediate use, but improved estimates can be obtained by iteratively looping over the observed light time series and identifying a stable model state that yields consistent phase markers, in this case, stable Dim Light Melatonin Onset (DLMO) predictions across successive iterations. This can be achieved by specifying the schedule and the number of iterations:

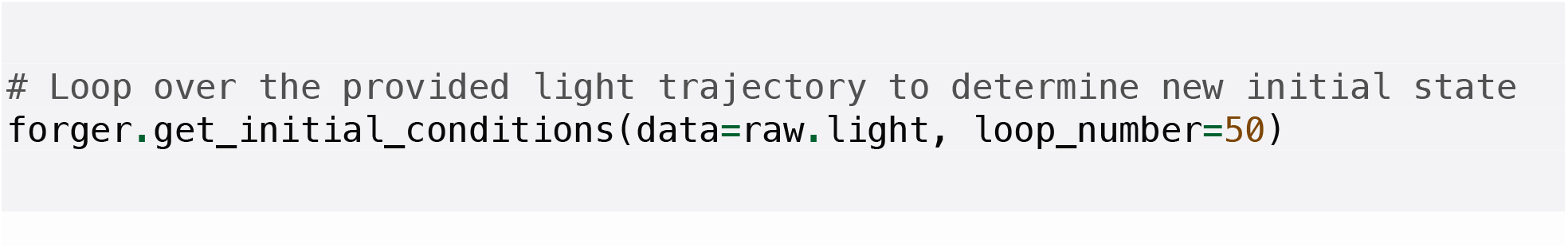

Daily predictions of DLMO can then be extracted as:

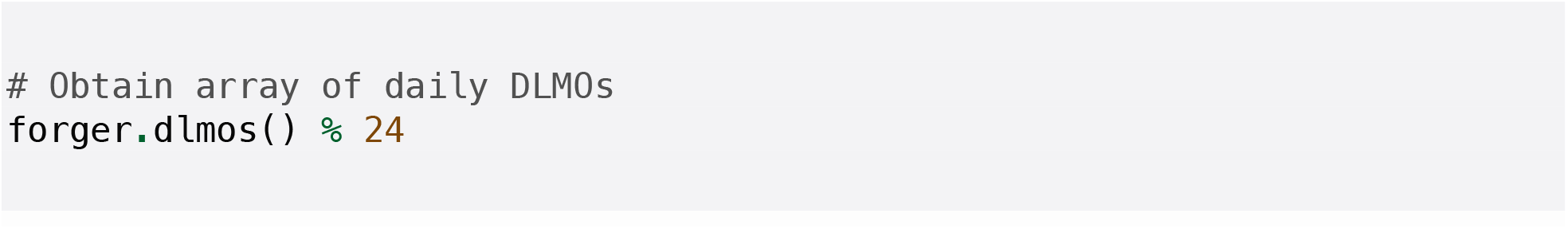

Model states can also be visualized using the `plot()` method:

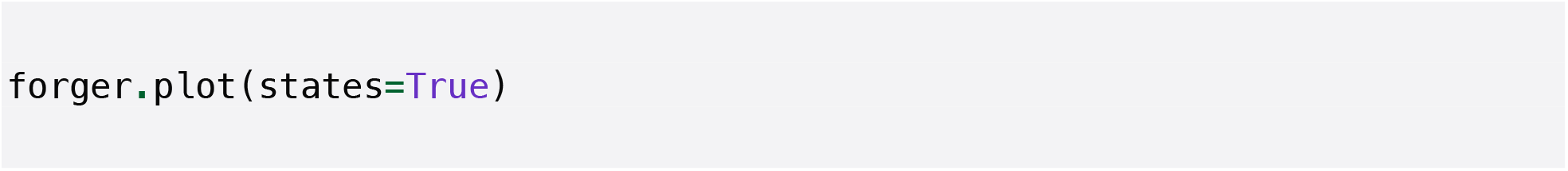

**Figure 2.**
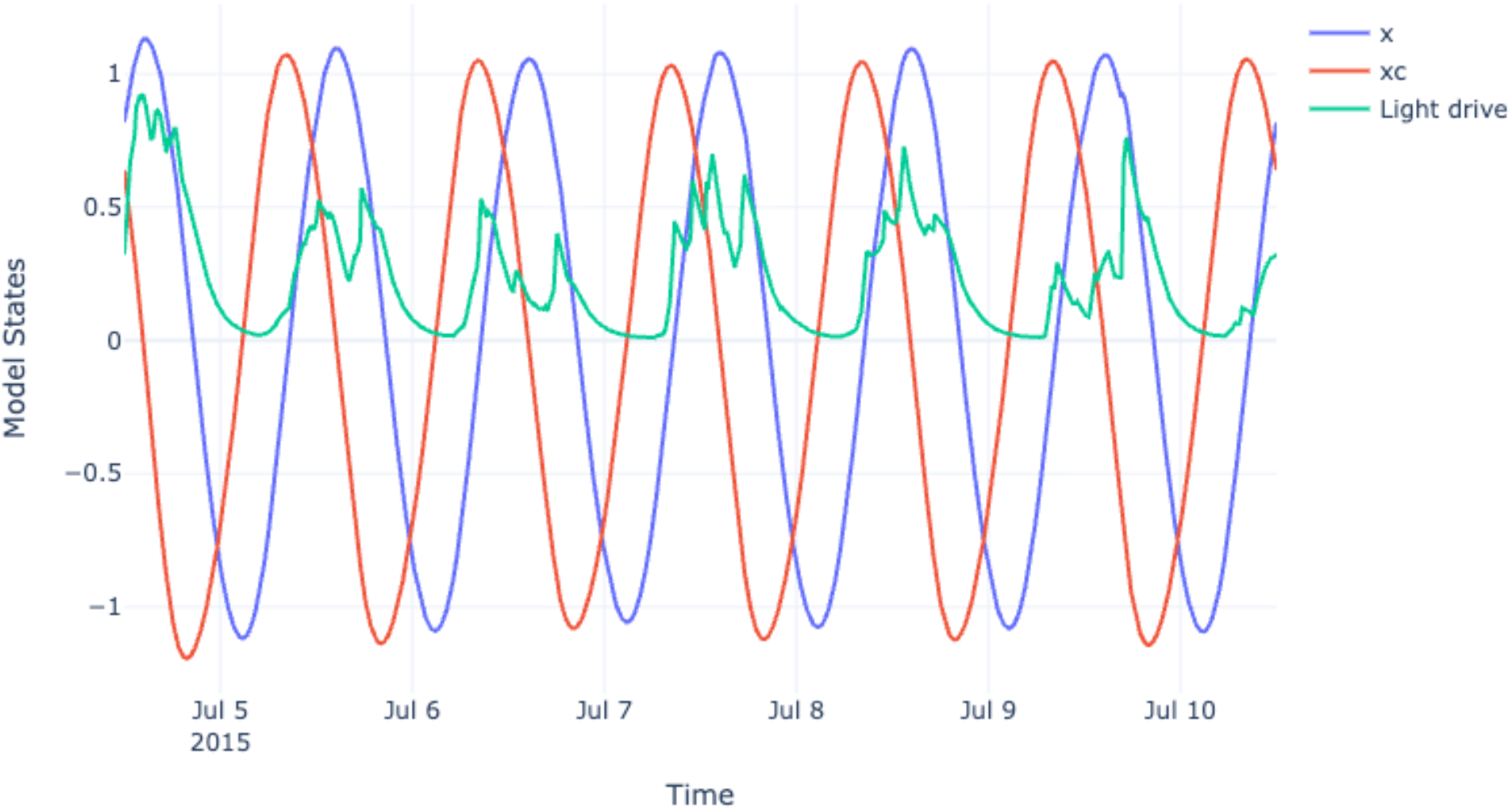
Interactive visualization of the three `Forger` model states (*x, x*_*c*_ and light drive) generated using the `plot()` method.

This is particularly useful for models whose state variables have physiological meaning, such as `breslow13`:

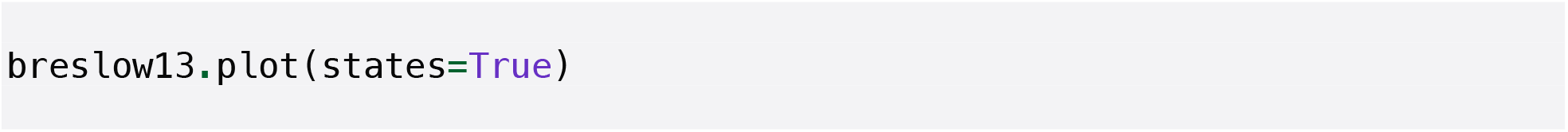

**Figure 3.**
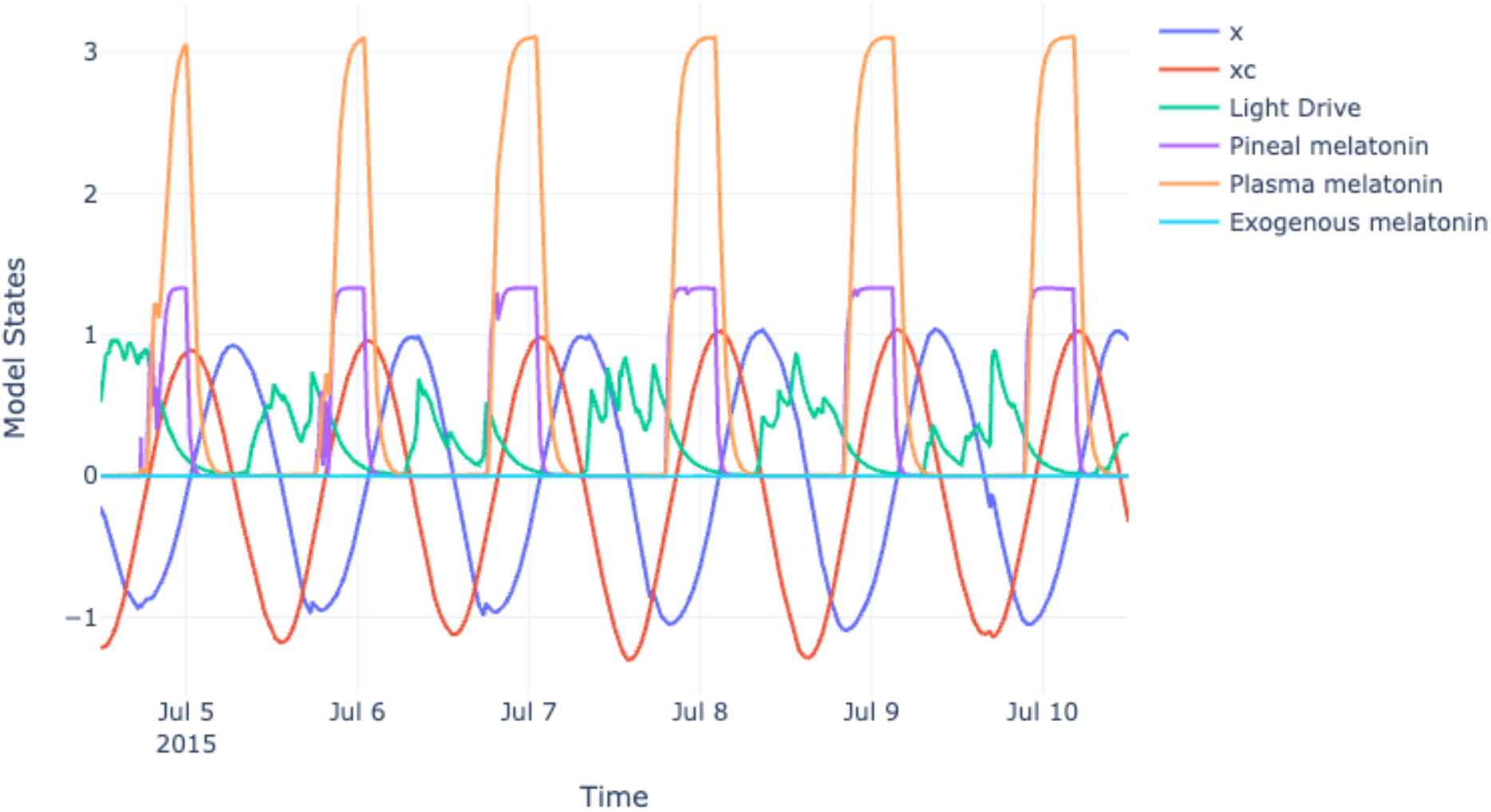
Interactive visualization of *Breslow13* model states. This model is able to predict the melatonin trajectory in three physiological compartments: pineal melatonin, plasma melatonin and exogenous melatonin.

### Predicting sleep pressure trajectory using the Skeldon23 model

The `Skeldon23` implementation follows a different workflow from the other models because it includes feedback between continuous and discrete components. Briefly, `Skeldon23` evolves four continuous state variables (*x, x*_*c*_, *n, h*) through numerical integration, while the sleep/wake state *s* is updated iteratively through a switching rule based on the current homeostatic and circadian state. In practice, the sleep state is held constant within each time interval, the continuous equations are integrated over that interval, and the sleep state is then updated at the interval boundary. Given the nature of this implementation, *circStudio* separates model instantiation from model execution in `Skeldon23`: creating the object defines parameters and inputs, whereas calling `run()` performs the numerical simulation and populates the predicted trajectories:

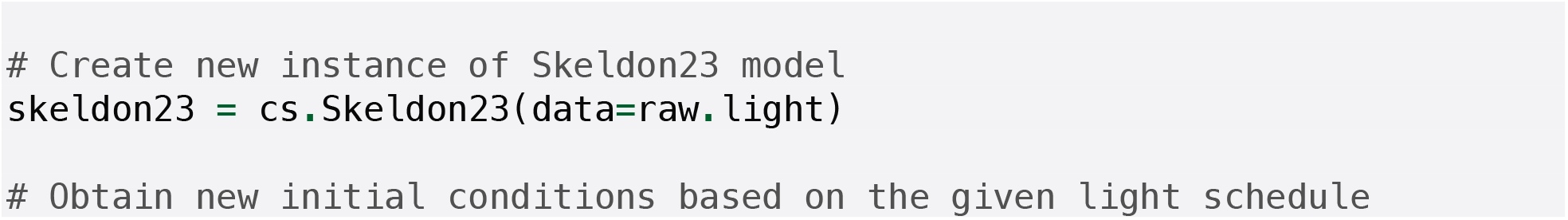

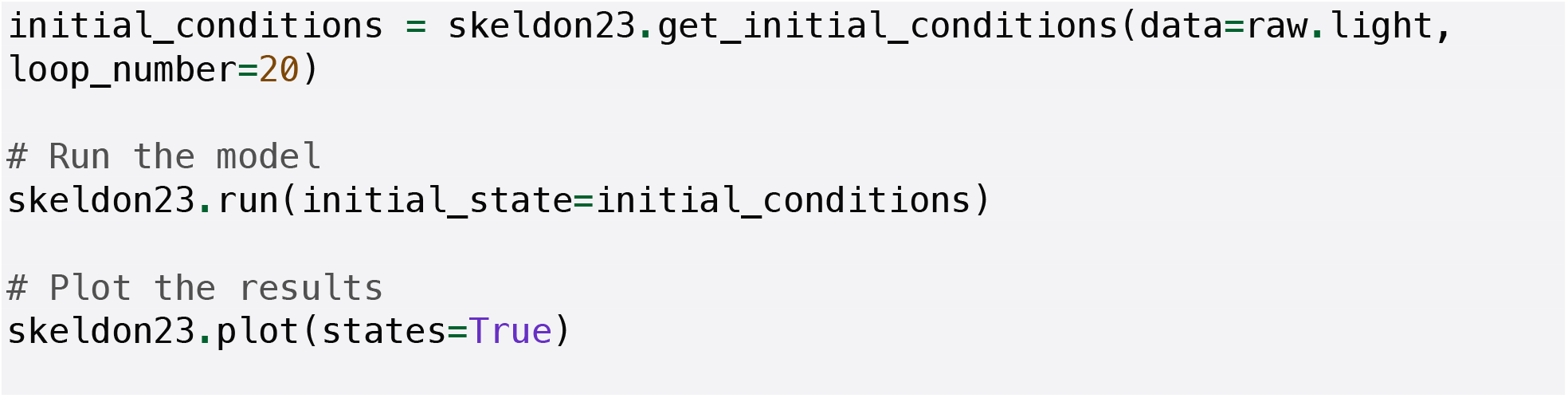

**Figure 4.**
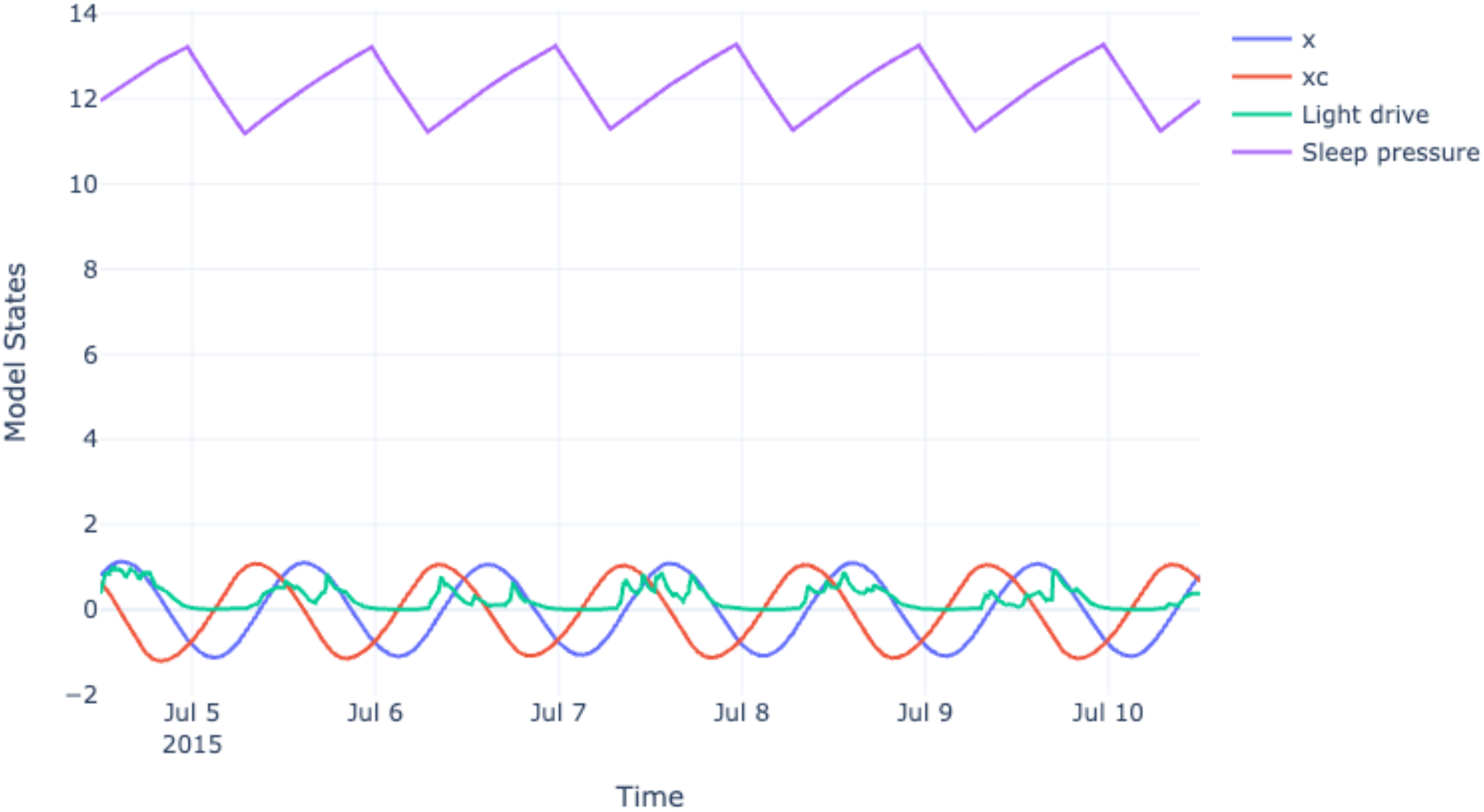
Visualization of model states of the *Skeldon23* model. This model is able to predict a trajectory corresponding to sleep pressure.

### Additional documentation and GitHub repository

In addition to the workflows described above, circStudio provides further functionality for Cosinor analysis, Singular Spectrum Analysis, (Multi-Fractal) Detrended Fluctuation Analysis and Functional Linear Modeling. These methods were migrated from *pyActigraphy* and improved documentation was written, including interactive examples:

**Table 1.** Online resources available for *circStudio*, including tutorials, documentation, and the source code repository.

| Resource | Link |
| --- | --- |
| Installation | https://djmarques.github.io/circStudio/tutorial_0.html |
| Loading actigraphy files | https://djmarques.github.io/circStudio/tutorial_1.html |
| Preprocessing actigraphy data | https://djmarques.github.io/circStudio/tutorial_2.html |
| Computing common metrics | https://djmarques.github.io/circStudio/tutorial_3.html |
| Mathematical models of circadian rhythms | https://djmarques.github.io/circStudio/tutorial_4.html |
| Cosinor analysis | https://djmarques.github.io/circStudio/tutorial_5.html |
| Singular spectrum analysis | https://djmarques.github.io/circStudio/tutorial_6.html |
| (Multi-fractal) Detrended Fluctuation Analysis | https://djmarques.github.io/circStudio/tutorial_7.html |
| Functional linear modelling | https://djmarques.github.io/circStudio/tutorial_8.html |
| API reference | https://djmarques.github.io/circStudio/api.html |
| Source code (GitHub repository) | https://github.com/djmarques/circStudio |

### Applying *circStudio* to a practical case: submarine study

To illustrate the use of circStudio, we considered a biological question derived from our previous work on a submarine mission (Marques et al., 2025). In that study, we identified the Sleep Regularity Index (SRI) and relative amplitude (RA) as candidate behavioral markers of shift-specific organization in a military submarine environment. Building on this, we asked whether reduced sleep regularity during time spent aboard the submarine was associated with lower relative amplitude. This hypothesis was motivated by the expectation that irregular sleep-wake patterns increase locomotor activity during the habitual sleep period, as measured by the average activity levels during the least five hours of the day, thereby reducing the contrast between rest and activity that underlies RA.

To answer this question, we first specify the directory containing the raw actigraphy files corresponding to the submarine mission period (here denoted `source`). We then iterate over all files in this directory:

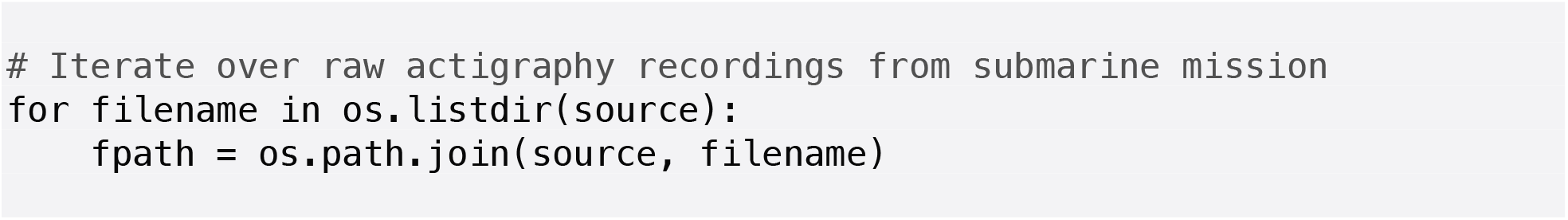

Within this loop, circadian metrics can be computed for each individual. It is possible to obtain RA, L5 and SRI as follows:

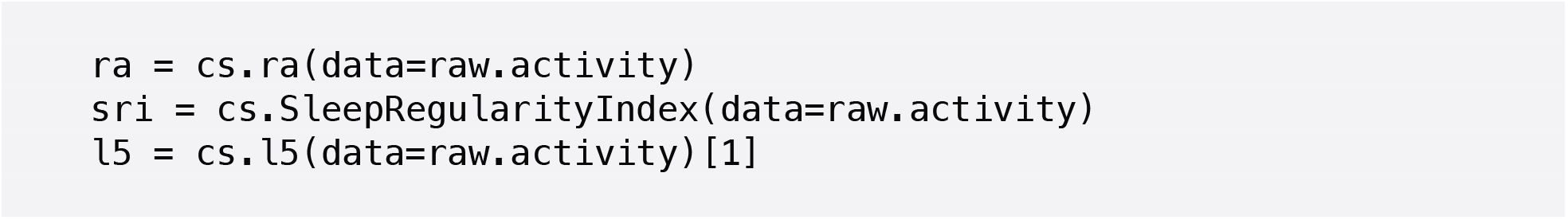

The resulting metrics were stored and converted into a tabular structure for downstream analysis. We then assessed the association between SRI and RA using Spearman’s rank correlation coefficient (SciPy), observing a moderate positive correlation (ρ = 0.422, p = 0.025). In addition, SRI was negatively correlated with L5 (ρ = −0.514, p = 0.025), consistent with increased activity during the least active period in individuals with more irregular sleep patterns. These findings are consistent with the hypothesis that reduced sleep regularity may be associated with diminished circadian rhythm robustness, potentially mediated by elevated nocturnal activity. However, given the observational nature of the analysis, these results should be interpreted cautiously and primarily as hypothesis-generating. This example illustrates how *circStudio* can be used to rapidly explore biologically meaningful relationships in actigraphy data and to guide the formulation of testable hypotheses in real-world settings.

## Discussion

A major contribution of *circStudio* is the provision of a unified platform for actigraphy analysis and data-driven simulation of circadian physiology within a single Python package. By combining flexible preprocessing tools, established actigraphy metrics, and several mathematical models of circadian rhythms, the package enables users to move efficiently from actimetry and wearable data to physiologically interpretable circadian outputs. This unification facilitates the development of health-oriented applications by providing direct access to research-grade analytical tools, while also offering a flexible foundation for building custom analysis pipelines for researchers in biological rhythms and neuroscience.

This integration of the actigraphy analysis capabilities of *pyActigraphy* with advanced circadian modeling tools from the *circadian* package developed by Arcascope is consistent with the original vision of *pyActigraphy*. Even though the *circStudio* codebase follows a different architecture, this redesign contributes to the broader goal of establishing a comprehensive toolkit for actigraphy analysis, and some functionalities developed in *circStudio* may be backported to *pyActigraphy* in future releases. Future *circStudio* versions will also introduce standard algorithms to automatically detect spurious periods of inactivity, thereby simplifying the identification and removal of non-wear periods.

One potential limitation of circStudio is its specialization in actigraphy analysis. A possible future direction would therefore be the development of a more general package for biological time series analysis, providing a broader toolkit applicable to diverse time-resolved datasets in biology. Additionally, even though the syntax is intentionally concise, the development of a graphical user interface could further broaden the accessibility of *circStudio* across circadian and sleep research.

